# *FT/FD-GRF5* repression loop directs growth to increase soybean yield

**DOI:** 10.1101/437053

**Authors:** Kun Xu, Xiao-Mei Zhang, Guolong Yu, Mingyang Lu, Chunyan Liu, Haifeng Chen, Jinlong Zhu, Fulu Chen, Zhiyuan Cheng, Penghui Huang, Xinan Zhou, Qingshan Chen, Xianzhong Feng, Yuchen Miao, Liangyu Liu, Iain Searle, Yong-Fu Fu

## Abstract

Major advances in crop yield are eternally needed to cope with population growth. To balance vegetative and reproductive growth plays an important role in agricultural yield. To extend vegetative phase can increase crop yield, however, this strategy risks loss of yield in the field as crops may not mature in time before winter come. Here, we identified a repression feedback loop between *GmFTL*/*GmFDL* and *GmGRF5-1* (*Glycine-max-Flowering-Locus-T/Glycine-max-FDL* and *Glycine-max-GROWTH-REGULATING-FACTOR5-1*), which functions as a pivotal regulator in balancing vegetative and reproductive phases in soybean. *GmFTL*/*GmFDL* and *GmGRF5-1* directly repress gene expression each other. Additionally, *GmGRF5-1* enhances vegetative growth by directly enhancing expression of photosynthesis- and auxin synthesis-related genes. To modulate the loop, such as fine-tuning *GmFTL* expression to trade-off vegetative and reproductive growth, increases substantially soybean yield in the field. Our findings not only uncover the mechanism balancing vegetative and reproductive growth, but open a new window to improve crop yield.

## Introduction

The global crop demand for human consumption and livestock feed is forecasted to increase by 110% from 2005 to 2050^1^, while recent advances on traditional breeding, genomics and transgenic technology are predicted to only yield improvements of up to 20%^2^. Flowering time is widely used as a selectable marker in plant high-yield breeding programs^3^, because vegetative growth is closely related to reproductive growth. A long vegetative phase means later flowering, larger and more vegetative organs (such as leaves and roots), that results in high yield because it provides the plenty source for yield formation. However, crops in the field may not mature normally before winter and lose the yield if the vegetative phase is too long^4^. Balancing vegetative and reproductive growth will achieve high yield in a normal growth season.

The transition from the vegetative to reproductive phase is regulated by a complex genetic network that monitors and integrates both the plant developmental and environmental signals, resulting in production of florigen *Flowering Locus T* (*FT*)^5,6^. The lower the florigen production, the later the flowering and the higher the yield, and *vice versa*^3^. The *FT* dosage plays a key role in the yield of tomato^7^ and rice^8^. Not only function on flowering control, *FT* homologs also contribute to the regulation of vegetative growth, such as tuberisation^9^, onion bulb formation^10^ and sugar beet growth^11^. *FT* coordinates reproductive and vegetative growth in perennial poplar^12^. Recently, *FT* was reported to regulate seed dormancy through *Flowering Locus C*^13^. *FT*, together with *FD* negatively correlates with leaf size as its overexpression leads to smaller leaves and reduced expression leads to increased leaf size^6,14^. But, the mechanism of *FT* in regulating the development and function of leaves is still uncovered.

*GROWTH-REGULATING-FACTOR* (*GRF*) family genes encode plant-specific transcriptional factors, which are shown to bind DNA with a TGTCAGG *cis*-element to repress or activate the expression of their target genes^15,16^. *GRF5* is confirmed as a key regulator of leaf growth and photosynthetic efficiency by enhancing cell division and chloroplast division, with concomitant increases in leaf size, longevity and photosynthesis^17^. Additonally, *GRF5* and cytokinins synergistically function in leaves^17^. Also, there are no direct target genes of *GRF5* in leaves identified yet.

Based on these observations, we hypothesised that *FT* participated in leaf development and then it is possible to gate *FT* expression to an extent that ensures that soybean to flower slightly later yet matures in the field before the low winter temperatures arrive, thereby increasing yield. In this study, we demonstrated that in soybean *GmFTL/GmFDL* directly inhibited *GmGRF5-1* expression; in turn, *GmGRF5-1* directly repressed *GmFTL* expression. Therefore, we established a repression feedback loop, which balanced vegetative growth and reproductive growth. Fine-tuning this loop by controlling *GmFTL* expression delayed flowering and increased yield in the field. *GmGRF5-1* directly activated genes related to energy metabolism (photosynthesis) and organ growth, which could all contribute to yield formation in soybean. Our results indicated that the *GmFTL/GmFDL*-*GmGRF5* repression loop was a new target to improve crop yield.

## Results

### Gating *GmFTL* expression increases yield

To test whether *GmFTL* controls soybean yield, we employed four selected transgenic lines of *GmFTL*-RNAi, which had lower expression of *Glycine max FT-like*s (*GmFTL*s) and later flowering^18^. Unsurprisingly, the soybean transgenic lines obviously showed high yield in both a growth chamber and greenhouse, and *GmFTL*-RNAi line 4 doubled yield per plant in the greenhouse (Fig. 1a, b). To obtain practical data in the field, we grew these soybean plants in two regions (Hanchuan and Beijing: N30°22’, E113°22’ and N39°58’, E116°20’, respectively). As a result of later flowering, only two lines (#1 and #3) matured normally in the growth season. These two lines significantly displayed higher yield, ranging from 14% to 62%, depending on the transgenic lines, locations and sowing time (Fig. 1c). TaqMan PCR data indicated that high yield related to lower expression of *GmFTL3*, a major florigen in soybean^19,20^ and that the level of *GmFTL3* transcripts was inversely proportional to yield (Fig. 1d). Together, florigen negatively contributed to plant yield.

**Fig. 1.**
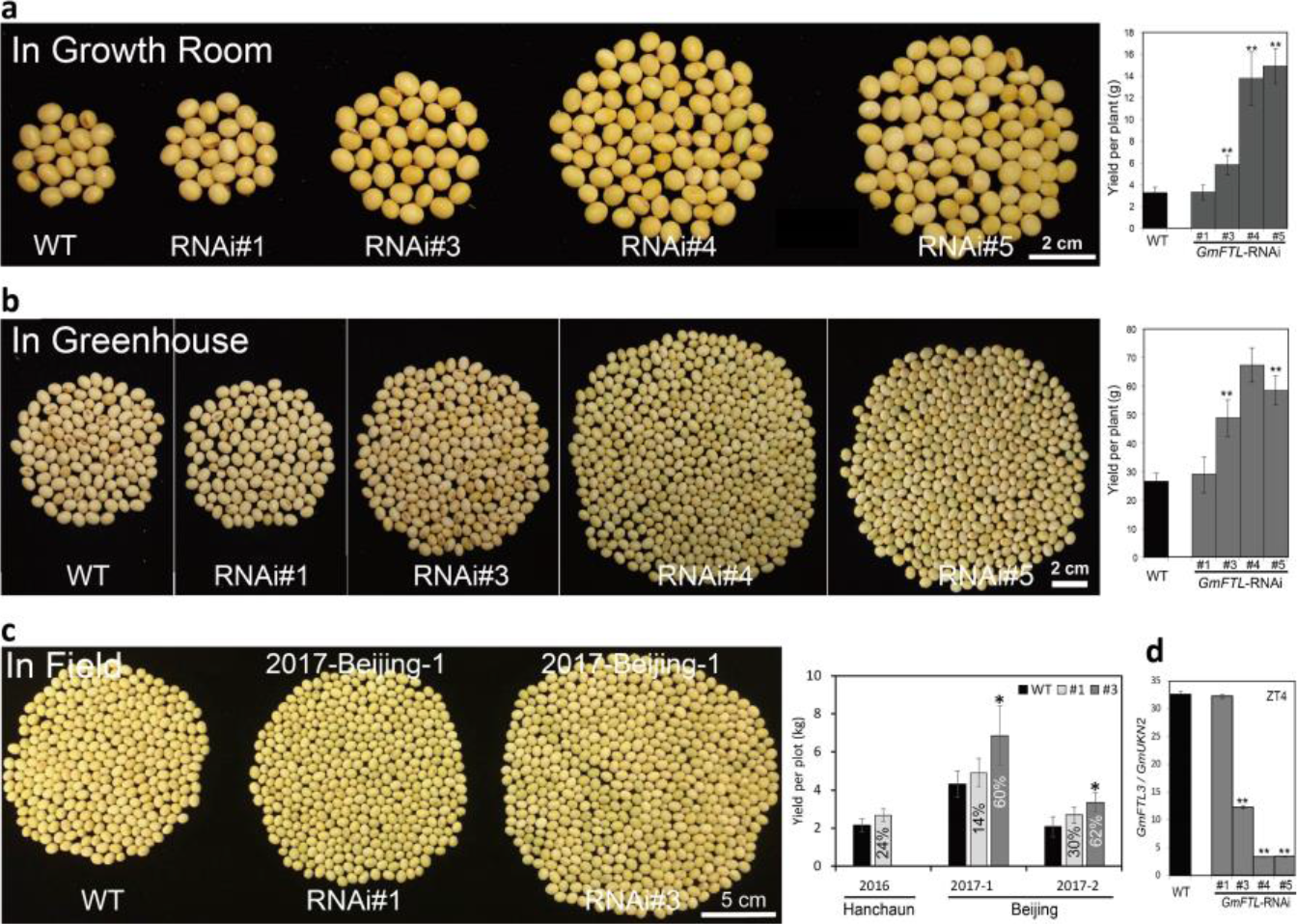
Reducing *GmFTL* expression increases soybean yield. **a**, **b** and **c**, Soybean yield in the growth room (**a**), glasshouse (**b**) and field (**c**). Left panels show the total number of seeds per plant. Right panels show the quantified yield per plant or plot. The percentages shown in the columns (**c**) indicate the increased yield compared to wild type. **d**, The mRNA abundance of *GmFTL3* in different transgenic lines. (* *P* < 0.05 and ** *P* < 0.01, Student’s *t*-test, n ≥ 10 plants).

### Yield increase contributed by photosynthesis

We next performed experiments to explain how *GmFTL* controlled soybean yield through growing *GmFTL*-RNAi lines in the greenhouse to analyse vegetative growth in detail. In the early stage of growth, there was little difference in the architecture between the transgenic and wild type plants. Along with growth, all RNAi lines displayed larger stature, with higher plants, more nodes, larger leaves and roots compared to WT, companying with later flowering and podding and high yield (Extended Data Fig.1, 2). Our results were consistent with a previous report on genome-wide association analysis for flowering and yield, which indicated a close correlation between plant height, flowering time and yield in soybean^21^.

Photosynthesis efficiency significantly affects plant yield^22^; thus, we wondered whether there was a change in the photosynthesis of the transgenic lines compared to the wild type plants. Transmission electron microscope revealed that the transgenic lines had a much more complicated structure in the chloroplast, with more and wider thylakoid membranes and rich grana in both the vegetative stage (the third trifoliolate opening) and flowering stage (the seventh trifoliolate opening) (Fig. 2a, Extended Data Fig. 3a). The detailed phenotypes are following (Fig. 2b, c, Extended Data Fig. 3b, c). Compared to wild type plants, the total number of chloroplasts (TNC) per cell of RNAi lines was lower, yet TNC per leaf area was higher in the vegetative stage because of the smaller cell size (CS). After flowering, the transgenic lines had larger CS and a lower TNC per leaf area. However, due to more and larger leaves, the transgenic lines had more TNC per plant at this stage. Transgenic cells also enriched different photosynthetic pigments (Fig. 2d).

Therefore, it was unsurprising that a biochemical assay evidenced that *GmFTL*-RNAi lines had much higher yield of the maximum quantum and photosynthetic rates (Figure 2d) and accumulated much more photosynthetic assimilates, such as starch, sucrose, glucose and fructose (Figure 2e). Thus, the transgenic *GmFTL*-RNAi plants had higher photosynthesis efficiency than did the wild type plants. Thus, the transgenic *GmFTL*-RNAi plants had higher photosynthesis efficiency than did wild type plants, which conferred to high yield as previous studies reported^22^.

**Fig. 2.**
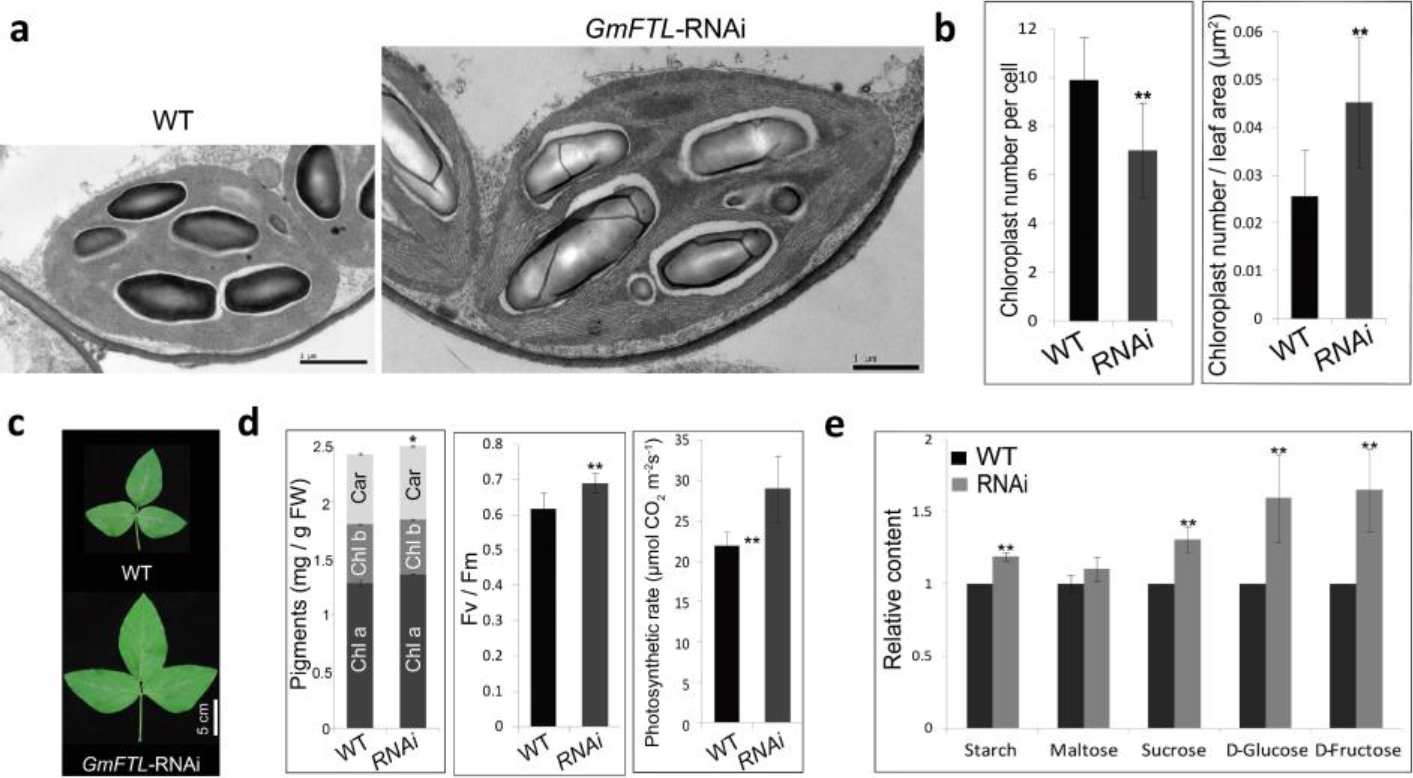
Reducing *GmFTL* expression enhances soybean leaf photosynthesis. **a**, The chloroplasts in the third trifoliolates. Scale bar, 1 μm. **b**, The average number of chloroplasts per cell and per leaf area in the third trifoliolates (n= 30 plants). **c**, The seventh trifoliolates. **d**, The amounts of chlorophyll a, b and carotenoids, the maximum quantum yield and the net photosynthesis (n= 5-10 plants). **e**, The contents of sugars in the 3^rd^ trifoliolates (n= 10 plants). Statistical analysis was done as Fig. 1.

### *GmFTL3/GmFDL* directly inhibits *GmGRF5-1* gene

During the floral transition, FT enhances flowering by interacting with FD in the shoot meristem, and FD binds to G-Box on its target genes^23^. *Arabidopsis GRF5* encodes a plant-specific transcription factor and regulates many plant-specific processes, such as chloroplast growth, photosynthesis and leaf growth^17^. The closest homolog of *GRF5* in soybean, *GmGRF5-1* (*Glycine max GRF5-1*), coded the protein shared highly conserved functional domains and localised in the nucleus (Extended Data Fig. 4a, b) as a transcriptional factor did. And in the promoter sequence of *GmGRF5-1*, there are two typical G-Box motifs, the binding site of bZIP transcription factor (such as FD) (Extended Data Fig. 5a). Together, we wondered if GmFDL may target directly *GmGRF5-1*, that in turn regulated photosynthesis.

Then we performed electrophoretic mobility shift assays (EMSA), and the results demonstrated that GmFDL5 (*Glycine max* FD-like5) proteins physically interact with G-Box I (Fig. 3a). Transcriptional activity analysis of transient expression in tobacco leaves indicated that *GmFDL5* inhibited *GmGRF5-1* promoter activity, and *GmFTL3*, the most likely candidate of florigen in soybean^19,20^, enhanced the effect of *GmFDL5* (Fig. 3b). Another FDL homolog, GmFDL1, also displayed inhibitory activity on the *GmGRF5-1* promoter (Fig. 3c). GmFTL3 proteins interacted with both GmFDL5 and GmFDL1 (Fig. 3d, Extended Data Fig. 6), that is consistent with a previous report^24^. To confirm the binding activity of *GmFDL in vivo*, we performed ChIP-qPCR with hair roots expressing *GmFDL5:GFP* and leaves of overexpressing *GmFDL1:GFP* plants, which showed early flowering (Fig. 3h). Both experiments identified that *GmFDL5*/*GmFDL1* enriched Fragment 4 of *GmGRF5-1* promoter (Fig. 3e). Further, the TaqMan PCR test exhibited that reducing *GmFTL* expression increased *GmGRF5-1* expression but had little effect on *GmFDL* expression (Fig. 3f). Therefore, we postulated that GmFDL functioned mainly with its binding activity, while GmFTL3 acted as a regulator and GmFTL3/GmFDL complex directly inhibits *GmGRF5-1* expression.

**Fig. 3.**
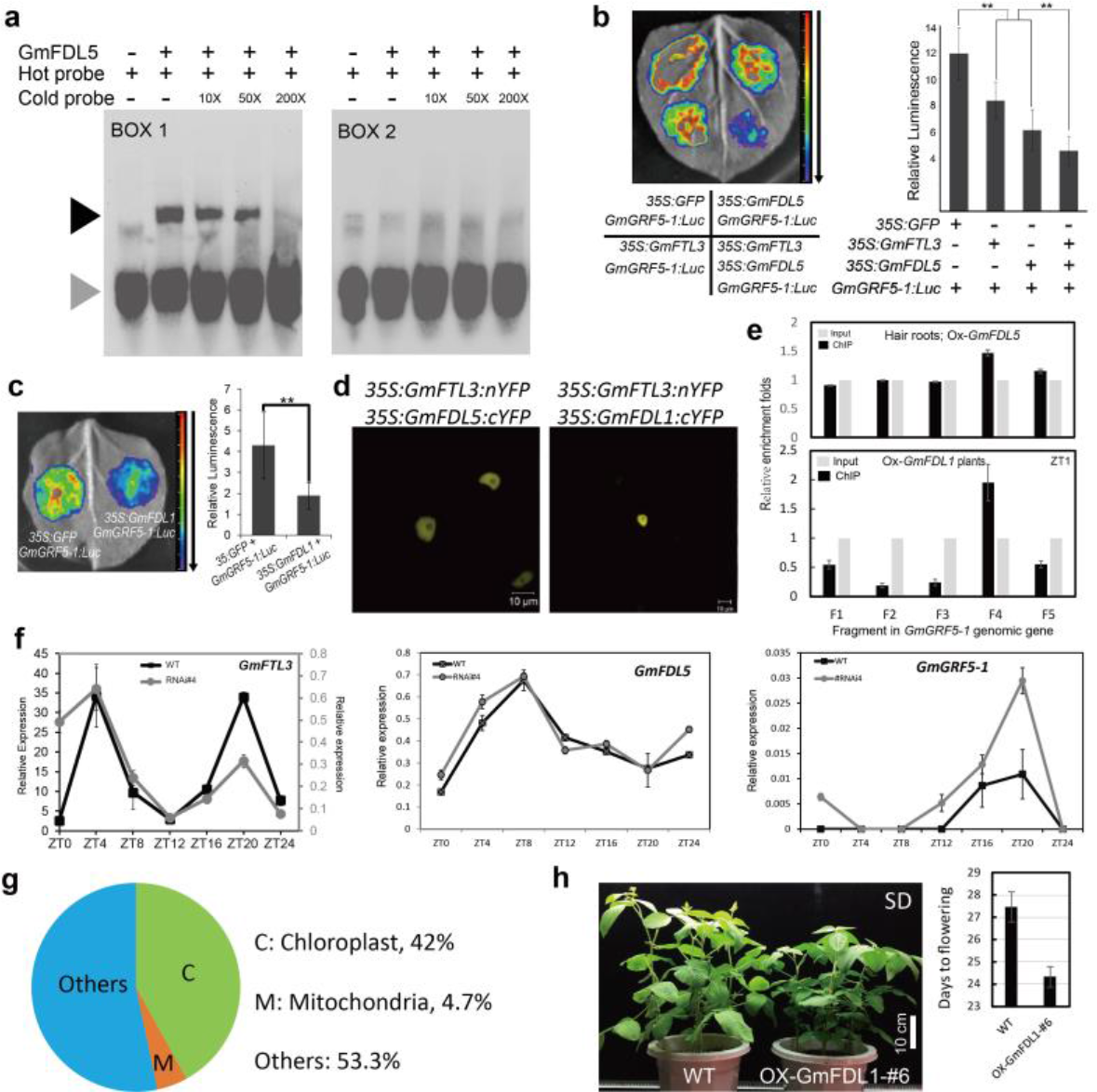
*GmFTL3/GmFDL* directly repress *GmGRF5-1* expression. **a**, EMSA: GmFDL5 bound to G-box 1 in *GmGRF5-1* promoter (Extended Data Fig. 5). **b**, **c,**Luciferase bioluminescence assay: *GmGRF5-1:Luc* co-expressed with *35S:GmFTL3* and *35S:GmFDL5* (**b**) or *35S:GmFDL1* (**c**). **d**, BiFC assay: GmFTL3 interacted with GmFDL1/5. **e**, ChIP-qPCR: FDL5:GFP enriched special fragments (relative to input) of *GmGRF5-1* promoter (Extended Data Fig. 5). **f**, RT-qPCR expression analysis of *GmFTL3*, *GmFDL5* and *GmGRF5-1*. **g**, RNA-seq: Genes with over two-fold change enriched to chloroplast (42%). **h**, Overexpressing *GmFDL1* soybean plants flowered earlier. (n=12). Statistical analysis was done as Fig. 1.

### *GmGRF5-1* directly enhances vegetative growth

To identify targets of *GmGRF5-1* controlling leaf growth and photosynthesis, we firstly carried out transcriptome analysis of *GmFTL*-RNAi leaves and found that only a small fraction of genes (0.378%, 212 out of 56,044 protein-coding loci) in the soybean genome (*Glycine max* Wm82.a2.v1, https://phytozome.jgi.doe.gov/) showed more than twofold changes in gene expression (Extend data Table 3), and RT-qPCR confirmed some of them (Extended Data Fig. 3e). Among these, 42% of genes encoded proteins targeted to the chloroplast and 4.7% of proteins targeted to the mitochondria (Fig. 3g), thereby suggesting that *GmFTL*-RNAi did have a significant effect on energy metabolism in soybean.

In *Arabidopsis*, GRF regulates target gene expression through direct binding to a TGTCAGG *cis*-element in target gene sequences^25^. We then randomly selected 10 genes from them as potential *<u>G</u>mGRF5-1* <u>t</u>arget <u>g</u>enes (*GTG*), due to their potential functions in photosynthesis (Extend data Table 3) and their sequences containing at least one TGTCAGG *cis*-element (Supplemental Information), a GRF binding sites^25^. These genes included hypothetical chloroplast open reading frame (*YCF1*, *GTG1* here), NAD(P)H dehydrogenase subunit H (*GTG2*), ATP synthase subunit alpha (*GTG3*), Plant basic secretory protein family protein (*GTG4*), DNA-directed RNA polymerase family protein (*GTG5*), Photosystem II reaction center protein D (*GTG6*), Photosynthetic electron transfer D (*GTG7*), Sucrose-proton symporter 2 (*GTG8*), Phosphoenolpyruvate carboxylase-related kinase 2 (*GTG9*), Nucleotide-sugar transporter family protein (*GTG10*). Surprisingly, *GmGRF5-1* significantly enhanced expressions of most of *GTG* genes in tobacco cells (Fig. 4a, c, Extended Data Fig. 7a). Then, we focused on *GTG1* gene, a member of *YCF* family with significant functions in gene expression and photosynthesis^26^. In *35S:GmGRF5-1:MYC* transgenic soybean plants, GmGRF5-1:MYC enriched Fragment 5 in the *GTG1* genome sequence (Fig. 4d, Extended Data Fig. 5b). GmGRF5-1 proteins physically interacted with this fragment in an EMSA analysis (Fig. 4e). What is more, in the presence of 35S-mini fragment, Fragment 4 and 5 of *GTG1* could replace the full length of *GTG1* promoter and be activated by GmGRF5-1 proteins (Fig. 4b). In *GmGRF5-1:MYC* plants, not only there was an increase of *GTG1* transcripts (Fig. 4f), but all of the leaf size, chlorophyll content, photosynthesis rate and complexity of chloroplast increased (Fig. 4g, h, i), that contributed to plant yield (Fig. 5a). Combining these results, we hypothesised that *GmFTL-*RNAi increased photosynthesis through at least enhancing the expression of *GTG* genes mediated by *GmGRF5-1*.

**Fig. 4.**
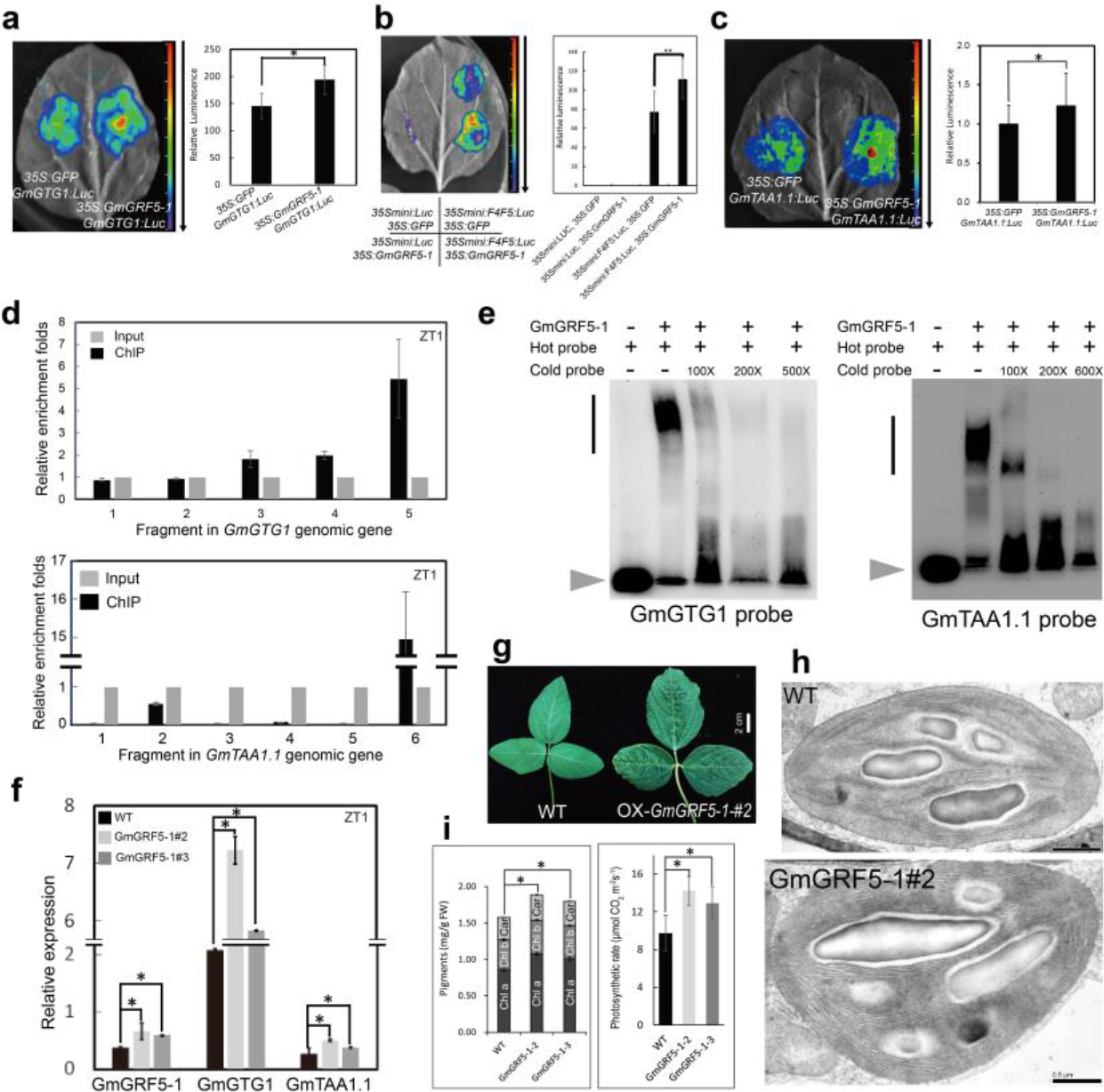
*GmGRF5-1* directly enhances *GmGTG1* and *GmTAA1.1* expression. **a**, **b**and **c**, Luciferase bioluminescence assay: *GmGTG1:Luc* (**a**), *35Smini:F4F5:Luc* (*F4* and *F5* fragments of *GTG1* (**b**) or *GmTAA1.1:Luc* (**c**) were activated by *35S:GmGRF5-1*. **d**, ChIP-qPCR: GmGRF5-1:MYC enriched special fragments (relative to input) of *GmGTG1* or *GmTAA1.1* genes. **e**, EMSA: *GmGRF5-1* directly bound to special fragments of *GmGTG1* and *GmTAA1.1*. **f**, RT-qPCR assay of *GmGRF5-1*, *GmGTG1* and *GmTAA1.1* mRNA. **g**, The third trifoliolates. **h,**Chloroplasts in the third trifoliolates. Scale bar, 1 μm. **i,**The amounts of chlorophyll a, b and carotenoids and the net photosynthesis (n= 10 plants). The fragments tested (**b**, **d**, and **e**) are shown in Extended Data Fig. 5. Statistical analysis was done as Fig. 1.

As a result of larger organs and stature in *GmFTL*-RNAi and overexpressing-*GmGRF5-1* plants, we supposed auxin may be involved in such phenotypes because auxin is the main phytohormone controlling organ growth and development^27^. *TAA1* is a key gene gating auxin biosynthesis in many plants^28^, and there are at least two *TAA1* homologs in the soybean genome (*GmTAA1.1-2*, Extended Data Fig. 7b). With similar strategy, we investigated the effect of *GmGRF5-1* on *GmTAA1.1* gene. All results from luciferase luminescence assay (Fig. 4c), ChIP-qPCR (Fig. 4d), EMSA analysis (Fig. 4e), and gene expression (Fig. 4f) supported that *GmGRF5-1* directly and positively regulated the expression of *GmTAA1.1* gene. And we also proved that a GRF-binding *cis*-elements (Supplemental Information) in 3’UTR of *GmTAA1.1* gene (Extended Data Fig. 5c) was a GmGRF5-1 targeting site. The results indicated that *GmGRF5-1* may participate in auxin signalling pathway by directly and positively regulating auxin biosynthesis.

### *GmGRF5-1* directly represses *GmFTL3* gene

The overexpressing *GmGRF5-1* lines also displayed late flowering and higher yield per plant (Fig. 5a), similar phenotypes of *GmFTL*-RNAi line (Fig. 1). By analysing the *GmFTL3* genomic region, we found several potential GRF-binding *cis*-elements (Supplemental Information). Therefore, we questioned whether *GmGRF5-1* could directly regulate *GmFTL* in a feedback mode, and we exploited the similar approaches above to address it. The first line of evidence from tobacco cells indicated that *GmGRF5-1* obviously inhibited expression of *GmFTL3* genomic gene (Fig. 5b), but not *GmFDL5* (Fig. 5c). ChIP-qPCR isolated *GmGRF5-1:MYC-* enriching Fragment 4 in *GmFTL3* genomic gene (Fig. 5e, Extended Data Fig. 5d), while EMSA provided a strong line of evidence for direct binding of *GmGRF5-1* to this fragment (Fig. 5d), which harboured a GRF *cis*-element (TGTCAGC) (Supplemental Information). Such an inhibitory effect of *GmGRF5-1* on *GmFTL3* expression was confirmed by RT-qPCR for the level of *GmFTL3* transcripts in the leaves of overexpressing *GmGRF5-1* lines (Fig. 5f). Thus, a late-flowering phenotype of overexpressing *GmGRF5-1* plants may result from a direct and negative effect of *GmGRF5-1* on *GmFTL3* expression.

**Fig. 5.**
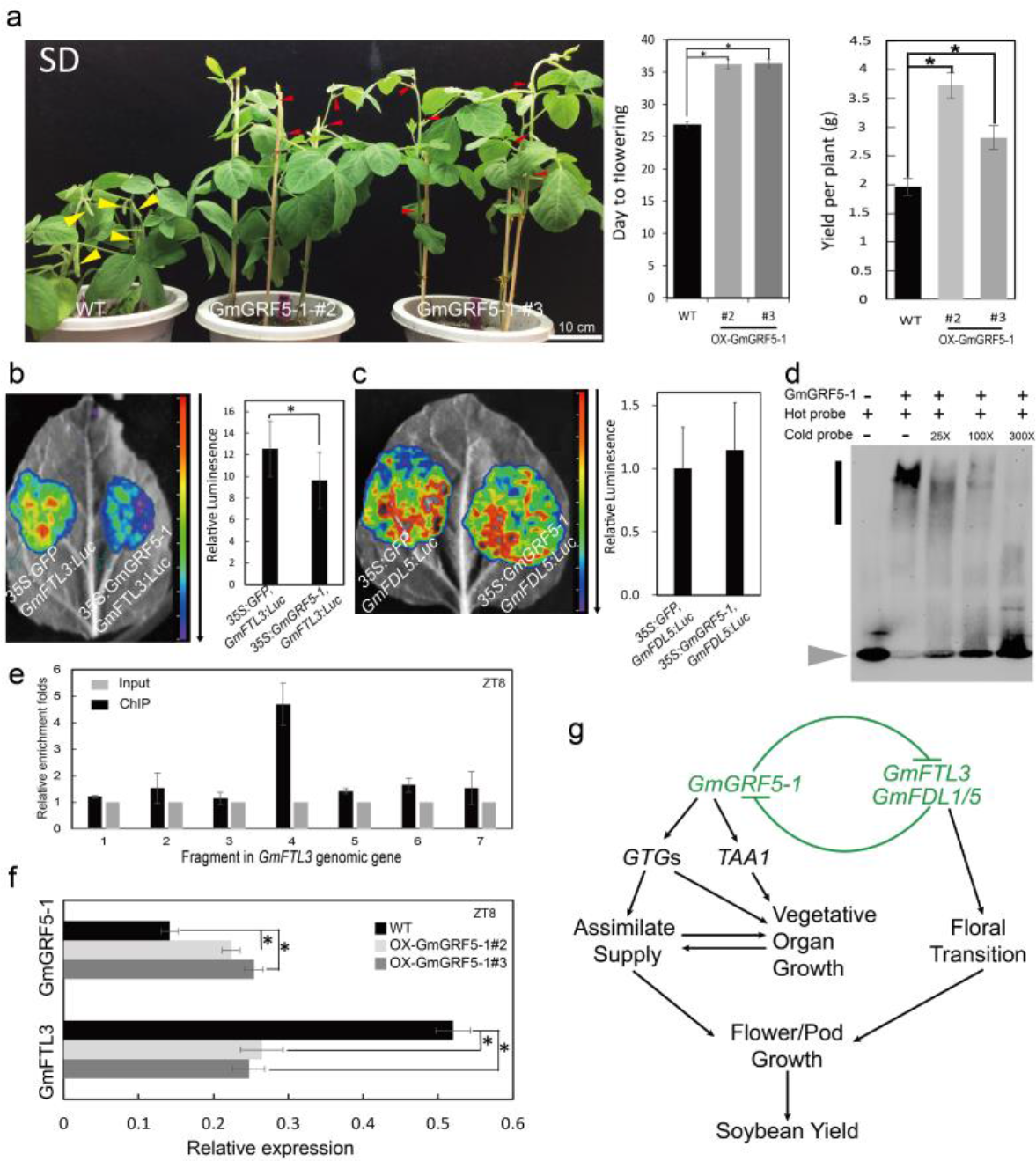
*GmGRF5-1* directly represses *GmFTL3* expression. **a**, Overexpression of *GmGRF5-1* delayed flowering and increased yield in soybean. Yellow arrows, developing pods; red arrows, flowers. Scale bar, 10 cm. **b**and **c**, Luciferase bioluminescence assay: *35S:GmGRF5-1* co-expressed with *GmFTL3:Luc* (**b**) or *GmFDL5:Luc* (**c**). **d.**EMSA: GmGRF5-1 binds to Fragment 4 of *GmFTL3* gene (Extended Data Fig. 5). **e**, ChIP-qPCR: GmGRF5-1:MYC enriched Fragment 4 (relative to input) in *GmFTL3* gene. **f**. RT-qPCR assay of *GmGRF5-1* and *GmFTL3* mRNA. **g**. A repression feedback loop model showing the interaction of *GmGRF5-1* and *GmFTL/GmFDL* balancing vegetative and reproductive growth. Statistical analysis was done as Fig. 1.

## Discussion

*FT* promotes flowering control^3,5,6^, while *GRF5* enhances chloroplast growth, photosynthesis and leaf growth^17^. Here, we identified a target gene of *FT* in leaves and a couple of target genes of *GRF5* in soybean. Beyond flowering control, *FT* also plays vital roles in other important processes, including the growth of tubers, bulbs and sugar beets, and seed dormancy^9-11,13^, even though leaf growth^6,14^. Regulation of *GmFTL* on *GmGRF5-1* enriches our knowledge about florigen functions in leaf physiology. On the other hand, *GmGRF5-1* feedback-represses *GmFTL3* expression. Based on our data, we propose a repression feedback loop between *GmFTL*/*GmFDL* and *GmGRF5-1*, which plays a key role in balancing vegetative growth and reproductive growth (Fig. 5g). *GmGRF5-1* enhances vegetative growth through two pathways: one is related to metabolism through directly activating expression of energy metabolism (photosynthesis) related genes including *YCF1* (*GTG1*, here)^26^; the other is involved in organ growth through directly enhancing TAA1 expression (auxin synthesis^28^). cytokinins is also associate with organ growth^17^, while leaf growth may be aided by retrograde signalling from the chloroplasts, as indicated in a previous study^29^, a couple of energy-related genes are positively regulated by *GmGRF5-1*. Positive interaction between the two pathways increases assimilate supply for reproductive growth (flower and pod/seed growth). In contrast, the *GmFTL*/*GmFDL* complex pushes plants going through floral transition and committing to flowering and pod/seed development, which is supported by assimilate supply from vegetative organs. Given that *GmGRF5-1* has no effect on *GmFDL* expression and that *GmFTL* enhances *GmFDL* inhibitory effect on *GmGRF5-1*, *GmFDL* functions through its binding activity to target DNA, as does *FD* in rice^23^, and contributes to regulation of *GmFTL* on *GmGRF5-1* expression. That means that the expression level of *GmFTL*, not *GmFDL*, mainly determines the activity of the *GmFTL*/*GmFDL* complex in *GmGRF5-1* regulation in soybean. The direction and intensity of plant development (vegetative or reproductive growth) are manipulated by the repression-loop balance between *GmGRF5-1* and *GmFTL*/*GmFDL*. Therefore, overexpressing-*GmGRF5-1* and *GmFTL*-RNAi soybean plants show similar phenotypes, such as late flowering and high yield. Such a repression feedback loop is commonly a regulatory motif for fine manipulation of various biological processes. For example, the molecular circadian clock is maintained by repression feedback loops in different organisms^30,31^. Because *FT* gene is a main integrator of multiple flowering pathways^3,5,6^, knocking-down of *GmFTL* expression affect only a small set of genes with significant expression change (0.378%, here) and *FT* mutation results in very late flowering and larger biomass^3,5,6^, it is reasonable to believe that the repression-loop identified here may play a key role in monitoring various environmental cues and buffering cell physiology to adapt different stresses for plants to fulfil the life cycle.

Increased crop yield is required to feed a growing population^1^, and researchers endeavour to improve crop yields through studying different potential metabolic targets^32^. Our results confirm that the repression feedback loop of *GmFTL*/*GmFDL* and *GmGRF5-1* can serve as a new target to improve crop yield. For example, it is possible to fine-tune the expression of *GmFTL* through RNAi approach to a level compatible to agricultural production to increase crop yield in field. In this study, we improved soybean yield as much as 62% higher yield in the field compared to wild type. *GmFTL*-RNAi line #3 may be a potential new elite of high yield for soybean breeding in future. Several studies have indicated positive dosage effects of the florigen gene *FT* on the yield of crops, including rice and tomato^7,8^. However, these reports are based on genetic strategy (heterosis), which may not be easy to be transformed into agricultural practice. It is credible to obtain much better elites with yield phenotype in the field if constructing and screening more *GmFTL*-RNAi transgenic lines. What is more, any cues controlling this repression feedback loop could be employed to modulate plant development and translated into new targets of high yield. Due to high conservation of *GmFTL*/*GmFDL* and *GmGRF5-1* genes across plants, the repression feedback loop may be found in different plant species and the strategy of high yield here could be transferred to other crops. To understand the function and regulation of the repression feedback loop in spatial and temporal mode will help us to uncover plant development mechanism in more detail and to exploit much more efficient strategy to increase crop yield.

## Additional information

**Extended Data Fig. 1** Agronomic traits of *GmFTL*-RNAi and wild type soybean plants

**Extended Data Fig. 2** Reduction of *GmFTL* mRNA promotes trifoliolate leaf growth

**Extended Data Fig. 3** Photosynthesis phenotype and transcriptome in *GmFTL*-RNAi line

**Extended Data Fig. 4** Alignment of GmGRF5-1 homologs and GmGRF5-1 protein subcellular localization

**Extended Data Fig. 5** Schematic representation of the genomic genes studied in this study

**Extended Data Fig. 6** GmFTL3 interacts in the nucleus with GmFDLs

**Extended Data Fig. 7** GmGRF5-1 enhances expression of genes related to photosynthesis

**Extended Data Table 1** Putative biochemical function and identities of genes in this study

**Extended Data Table 2** List of oligonucleotide and primer sequences in this study

**Extended Data Table 3** Transcriptome analysis of GmFTL-RNAi line relative to wild type

**Supplemental Information** The sequences of genes and promoters studied in this study

## Acknowledgements

We wish to thank all friends and colleagues who helped us in any way but are not included in the author list. This research was supported by National Key R&D Project (Grant Nos. 2016YFD0101900 and 2016YFD0101005) from the Ministry of Science and Technology of China, the National Transgenic Research Project (Grant No. 2016ZX08004-005), the National Natural Science Foundation of China (Grant Nos. 31771714, 31371703 and 31570289), the CAAS-Innovation Team Project and the Basal Research Fund of CAAS (Y2017CG25), and partially funded by an Australia-China Science and Research Fund grant ACSRF48187 awarded to I.S.

## Author contributions

Conceptualisation: Y.-F.F. and X.-M.Z.; Methodology: K.X., Y.-F.F., G.Y., X.-M.Z., M.L., C.L. and H.C.; Investigation: K.X., X.-M.Z., G.Y., M.L., C.L., H.C., Y.-F.F., J. Z., F.C., Z.C., P.H, Y.M. and L.L.; Formal analysis and validation: K.X., X.-M.Z., G.Y., I.S., M.L. and Y.-F.F.; Visualisation: K.X. and Y.-F.F.; Writing—original draft: Y.-F.F. and K.X.; Writing—review and editing: I.S. and Y.-F.F.; Funding acquisition: X.F., X.-M.Z. and Y.-F.F.; Project administration: Y.-F.F. and X.-M.Z.; Resources: X.Z, Y.-F.F., Y.M., Q.C and I.S.; Supervision: Y.-F.F.

## METHODS

### Plant materials and growth conditions

*GmFTL*-RNAi plants were previously generated in the soybean (*Glycine max* (L.) Mer.) cultivar Tianlong1 in our lab^18^. All RNAi lines used here are independent, homozygous, transgenic lines. Wild type soybean controls for all experiments was cultivar Tianlong1. All plants were grown in controlled temperature and photoperiod growth rooms, greenhouses and/or the field. The light conditions in the plant growth rooms were short day conditions (8 hr light / 16 hr dark) from a LED light source with intensities ranging from 500 μmol·m-1·s-1 at soil level to 800 μmol·m-1·s-1 at the top of plants at maturity. Soybean field (plot) experiments were carried out at Hanchun (N30°22’, E113°22’) and Beijing (N39°58’, E116°20’). Spring sown soybeans were planted on 23rd April 2016 for Hanchun, and 15th May 15 or 1st June 2017 for Beijing. The plot area was 300 × 225 cm^2^ with 45 cm of row-spacing and 20 cm of plant-spacing at Hanchun or 600 × 300 cm^2^ with 60 cm of row-spacing and 30 cm of plant-spacing at Beijing. Agronomic characters were observed at maturity. Individual plants from each plot were subjected to statistical analysis.

### Plasmid construction and plant transformation

The encoding full-length sequence of *GmGRF5-1* was cloned into the binary vector pFGC5941-GW, a modified vector from pFGC5941, by an LR reaction to generate binary vector *35S*:*GmGRF5-1*:*MYC*, which was introduced into soybean cultivar Tianlong1 through cotyledon node transformation mediated by *Agrobacterium tumefaciens*. Transgenic lines were selected by Basta^®^ herbicidie resistance and transgene expression was measured by RT-qPCR.

To express His-tagged proteins in *E. coli*, *GmFDL5* and *GmGRF5-1* genes were cloned into pET28a (Novagen) with *Xba* I/*Bam*H I or *Nde* I/*Xho* I restriction enzyme sites, respectively. For *GmGRF5-1* promoter activity and ChIP analysis, the coding sequences of *GmFTL3*, *GmFDL5* and *GmFDL1* and the promoter sequence of *GmGRF5-1* were cloned into pGWC33, respectively. Then, pGWC-*GmFTL3*, pGWC-*GmFDL5* and pGWC-*GmFDL1* genes were LR-reacted with pGWB5 by LR reactions (Invitrogen) to get binary vectors of pGWB5-*GmFTL3*, pGWB5-*GmFDL5* and pGWB5-*GmFDL1*. pGWC-*GmGRF5-1* was combined with pGWB35 by LR reactions (Invitrogen) to get plant expressing vector of pGWB35-*GmGRF5-1*. For GmFDL5 and GmFDL1 ChIP experiment, the pGWB5-*GmFDL5* was used to root hair transformation, while pGWB5-*GmFDL1* vectors was employed for soybean stable transformation.

For other promoter analyses, the promoter and fragment sequences of interest were cloned into entry clone Fu76 (this lab) recombined into pSoy2 (this lab) by LR reactions (Invitrogen) to generate pSoy2-Luc destination clones34. pGWB6 was used to express GFP as a negative control. The promoter and fragment sequences were listed in Supplemental Information.

For BiFC assays, *GmFTL3*, *GmFDL5* and *GmFDL1*genes were cloned into pGWC. Then, by LR reactions (Invitrogen), the full length *GmFTL3* gene was cloned into pEarlyGate201 to generate a *GmFTL3-nYFP* binary vector, while full length *GmFDL5* and *GmFDL1* genes were cloned into pEarlyGate202 recombined to create *GmFDL5-cYFP* or *GmFDL1-cYFP* binary vectors.

*GmGRF5-1* gene was cloned into Fu2834 and the subsequent GFP-tagged gene pSoy2-*GmGRF5-1* was used for cellular localization analysis.

All PCR amplified genes used in this study were confirmed by Sanger sequencing. The sequences of the genes and promoters are listed in Supplemental Information. The description of genes and primer sequences for genes are listed in Extend data Table 1 and 2, respectively.

### RNA extraction and expression analysis

For RT-qPCR analysis, total RNA was extracted using EasyPure^®^ RNA Kit (ER101-01, TransGen Biotech). The quantity was measured by Nanodrop 2000C (ThermoFisher Scientific). For SYBR detection of RT-qPCR products, 500-1000 ng of the total RNA was used for reverse transcription (KR106-02, TIANGEN). SYBR Premix Ex-Taq (Perfect Real Time; TAKaRa) was used for the RT-qPCR assays. For TaqMan analysis, about 1 µg of total RNA was used for reverse transcription (KR106-02, TIANGEN), TaqMan™ Gene Expression Master Mix (No.4369016, ThermoFisher Scientific) was used for the RT-qPCR assays. The RT-qPCR was conducted using StepOne Plus (ABI). The reference genes *GmACT11* or *GmUKN2* were used as internal controls for normalization of all SYBR RT-qPCR data. Sequences of the primers are listed in Extend data Table 2. The 2-∆CT method was used to calculate the relative expression levels based on three technical replicates.

### Electrophoretic mobility shift assay (EMSA)

Recombinant proteins were purified from *E. coli* BL21 cells using nickel beads (No.31014, QIAGEN) for His-GmFDL5 or His-magnetic beads (Beaver Company, China) for His-GmGRF5-1 and purity confirmed by western blotting using an anti-His antibody (AbMart company, China). The purified proteins were dialyzed against Tris-HCl buffers (25 mM, pH 7.5) to remove contaminating imidazole. Biotin was used to label the 5’ end of both sense and antisense oligo probes. Probes were heated and cooled to obtain probe duplexes. The biotin-labeled DNA was synthesized by Invitrogen (Shanghai, China) for *GmGRF5-1* and BGI (China) for *GmFTL3* and *GmGTG1*, respectively. EMSA was carried out according to the manufacturer’s protocol for Light Shift Chemiluminescent EMSA kit (No. 20148, ThermoFisher Scientific). The binding buffer for *GmFDL5* protein binding assays contained 10 mM Tris-HCl (pH 7.5), 50 mM KCl, 1 mM DTT, 2.5% glycerol, 5 mM MgCl_2_, 50 g·μl-1 poly(dI·dC), 0.05% NP-40, while the binding buffer for GmGRF5-1 protein binding assay consisted of 10 mM Tris-HCl (pH 8.7), 50 mM KCl, 5 mM MgCl2, 10 mM DTT, 2.5% glycerol, 20 mM EDTA. Each 20 μl binding reaction contained 1 fmol of biotin-labeled, unlabeled dsDNA at different concentrations and recombinant His-tag proteins. Binding reactions were incubated for 30 min at 22°C. The reactions were separated on a 5% polyacrylamide gel for GmFDL5/GmGRF5-1 complex, or an 8% precast SurePageTM Gel (GenScript, China) for GmGRF5-1/GmFTL3.F4, GmGRF5-1/GmGTG1.F5 and GmGRF5-1/GmTAA1.1F6 complex binding assays. The running buffer used was TBE (pH 8.0) for GmFTL5 proteins or TBE (pH 8.75) for GmGRF5-1 proteins. Detection of biotin-labeled DNA was performed by chemiluminescence using a ChemiDoc-It Imaging System (UVP, Cambridge, UK).

### Bimolecular fluorescence complementation (BiFC)

The pEarlyGate201-*GmFTL3-nYFP* and pEarlyGate202-*GmFDL5-cYFP* or pEarlyGate202-*GmFDL1-cYFP* binary vectors were transiently expressed using *Agrobacterium tumefaciens*. Both recombinant *Agrobacterium* cells were co-infiltrated into *Nicotiana benthamiana* leaves. Empty vectors were used as negative controls and AtAHL22-RFP was used as a nuclear marker. *N. benthamiana* was grown under long-day (16 hr light:8 hr dark) conditions at 22℃ for at least 48 hours post infiltration. Leaves were observed under a confocal microscope (Zeiss LSM700).

### Luciferase bioluminescence assays

Luciferase assays were performed as previously outlined with some minor modifications35. Briefly, *A. tumefaciens* containing different vectors were co-infiltrated into 4-week-old *N. benthamiana* leaves. After co-infiltration, the plants were then grown in darkness for 12 hours and subsequently grown under long-day conditions (16hr Light:8hr Dark) for 1 day. Prior to bioluminescence imaging, co-infiltrated leaves were uniformly sprayed with 2 mM D-luciferin (Gold Biotechnology) diluted in 0.01% Triton X-100 solution. After spraying with D-luciferin, leaf samples were incubated in the dark for 5 min. Exposure times for luminescence imaging were 3-5 min. Imaging of bioluminescence signals were performed using a NightShade LB 985 *In vivo* Plant Imaging System (Berthold Technologies, Germany). IndiGO Imaging Software (Berthold Technologies, Germany) was used for image acquisition and processing of luminescence. At least ten leaves for each experiment were used, and each experiment was repeated at least three times. All potential promoter sequences of genes except *GmFTL3* and *GmTAA1.1* were employed in this study. The full sequence of promoter, 5’UTR, genomic gene and 3’UTR of *GmFTL3* (Figure 5D) was used for this assay and the luciferase gene was inserted just before the stop codon of *GmFTL3* gene. The sequence of promoter and 3’UTR of *GmTAA1.1* was used to analyze *GmGRF5-1* activity on Gm*TAA1.1* regulation. All promoter sequences fused to luciferase are listed in Supplemental Information.

### Chromatin immunoprecipitation (ChIP) qPCR analyses

Chromatin was extracted from leaves or roots of soybean plants grown under short day conditions. Tissue was harvested at zeitgeber 1 (ZT1) timepoint. ChIP-qPCR assays were performed essentially as described36 with minor modifications: (1) cross-linking duration was 30 min with three intervals for stirring the solution to remove bubbles on the surface of leaves, (2) sonication time was 20 min, (3) two steps were employed to reverse cross-link, that included NaCl (300 mM) incubation overnight at 65°C and 1% Chelex-100 for 15 min at 95°C. The sonicated DNA were used for immunoprecipitation with commercially available anti-GFP-mAb-magnetic beads (MBL, D153-11) for the GmFDL5-GFP assay and anti-Myc-mAb-magnetic agarose (MBL, Mo47-10) for GmGRF5-1 assay. After reversing the cross-linking, immunoprecipitated DNA was analyzed by qPCR using primers (Supplemental Table S1) for specific regions. For the GmGRF5-1 binding assay, the regions for qPCR analysis contained at least one of the AtGRF7 protein binding motifs (TGTCAGG25) or its derivatives (TGTCAAG, TGTGAAG, TGTTAGG, TTTCAGG, GATCAGG, GTTCAGG, TGTCATG, TGGCAGG, CTTCAGG, TGTCACG TGTTATG). Three independent experiments, each using around 1 g of leaves were performed. Three technical replicates for each qPCR were carried out and Gm*ACT11* or *GmUKN2* were used as internal controls for normalization. Each experiment was repeated at least three times.

### Chloroplast analysis and chlorophyll measurements

The fully-opened third and seventh trifoliolate leaves of soil-grown greenhouse WT (cv. Tianlong1) and *GmFTL*-RNAi line 4 soybean plants were harvested for the measurements. Perpendicular transverse sections of the middle leaflets of trifoliolate leaves were prepared by the Transmission-Electron-Microscope and Mass-Spectrometry Platform of Institute of agricultural products processing, Chinese Academy of Agricultural Sciences (Beijing, China). The photos were obtained by using a transmission electron microscope (H-7500, Japan).

Microscopic differential interference contrast images were taken, the area of 200 mesophyll cells flanking the epidermis was measured by using ImageJ software. Chloroplast number per leaf area and chloroplast number per cell were measured by using Image J software.

For chlorophyll content analysis, we punched 20 fresh sections (d = 6 mm) from five individual leaves for each sample by using a hole puncher. The samples were immersed in 25 mL of 80% acetone, and stored at room temperature for 5 days. Then 1 mL of the supernatants was measured for absorbance at 663 and 645 nm. The concentration of chlorophyll a and b (Chl a and Chl b) and carotenoids was calculated using the following formulas:

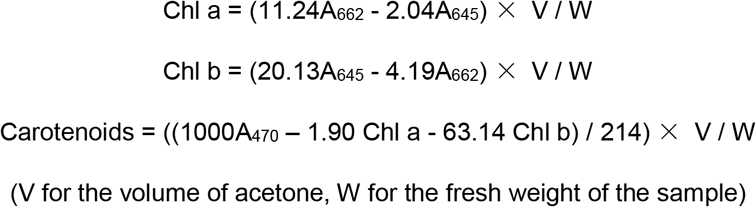

### Photosynthetic rate and chlorophyll fluorescence analyses

Plants were grown under SD conditions, and the third trifoliolate leaves (n = 10) of WT (cv. Tianlong1) and *GmFTL*-RNAi line 4 line were selected for measuring the photosynthetic rates at days 28 to 35 after sowing following the manufacturer’s instructions (LICOR LI-6400 V4.0.1). Fv/Fm was measured using an IMAGING-PAM M series Chlorophyll Fluorometer with the MAXI version (Heinz Walz, Effeltrich, Germany). SD grown plants were placed in darkness for 30 min before measurements were taken.

### Measurement of sugar content in leaves

WT (cv. Tianlong1) and *GmFTL*-RNAi line 4 seeds were sown in spring 2015 and grown at Beijing (China). Third fully-opened trifoliolate leaves were harvested at one hour after sunset for the measurement of sugar contents (n = 5). For the extraction of soluble sugars and starch, about 100 mg of leaf samples were homogenized in 0.5 mL of 80% (v/v) ethanol in a 1.5-mL tube and incubated at 70°C for 90 min. Following centrifugation at 16,000 g for 5 min, the supernatant was transferred to a new 1.5 mL tube. The pellet was rinsed twice with 2 mL of 80% ethanol and removed. Any remaining solvent was evaporated at room temperature by using a vacuum. The residue was resuspended in 0.3 mL of distilled, sterile water and this represented the soluble carbohydrate fraction. The remaining pellet contained the insoluble carbohydrates, included starch, was homogenized in 0.2 mL of 0.2 N KOH and the suspension was incubated at 95°C for 1 hr to dissolve the starch. Following the addition of 0.035 mL of 1 N acetic acid and centrifugation for 5 min at 16,000 g, the supernatant was used for starch quantification. Detailed procedures were followed according to the manufacturer’s instructions; starch (No. 1013910603), Maltose/Sucrose/D-Glucose (No. 11113950035) and D-Glucose/D-Fructose kits (No. 10139106035) (R-Biopharm, Germany).

### Measurement of leaflet area

When the trifoliolates were fully opened (plants were grown in greenhouse), the length and width of each leaflet were measured. And so on, for each of trifoliolates of indicated lines. Leaflet area was calculated using the formula d = (L + 2W) / 3 (d indicates the diameter; L represents the length of leaflet; W represents the width of leaflet). (n = 7 plants)

### Leaf anatomical structure analysis

Fully-opened third or seventh trifoliolate leaves of soil-grown, glasshouse plants of WT or *GmFTL*-RNAi line 4 plants were excised. The leaves were fixed in FAA fixative solution (formaldehyde solution 5 ml: glacial acetic acid 5 ml: 70% ethanol 90 mL). Perpendicular transverse sections were prepared through the middle leaflets of trifoliolate leaves by Servicebio Biotechnology Co Ltd (Wuhan, China). The sections were observed, and photographs were obtained by using a SteREO Discovery V20 (Zeiss, Germany).

### Measurement of major agronomic traits

The flowering time was determined by emergence of the first flower on the main stem of soybean plants. Podding time was determined as the time when the first pod on the main stem was 2 cm in length. Measurements of plant height, branch number, node number, pod number and seed number per plant were performed at full plant maturity. For experiments in both the greenhouse and growth rooms, more than ten individual plants of each line were sampled for analysis of all those traits. To examine field yield traits, three plot replications were used at Beijing (N39°58’, E116°20’) and Hanchuan (N30°63’, E113°591) sites for each indicated line. For the Beijing field experimental site, the planting density was 30 cm × 60 cm, with one plant per site. The area per plot was 14.4 m2. For the Hanchuan plot site, the planting density was 20 cm × 50 cm, with two plants per site. The area per plot was 6 m2.

### Measurement of leaflet area

When the first trifoliolate leaves were fully opened, the length and width of each leaflet of plants (n = 7 plants) grown in the greenhouse were measured. Leaflet area was calculated using the formula d = (L + 2W) / 3 (where d indicates the diameter; L represents the length of the leaflet; W represents the width of the leaflet).

### Transcriptome analysis

The fully-opened third trifoliolate leaves from WT or *GmFTL*-RNAi line 4 line plants were sampled one hour after sunrise. Total RNA was extracted using EasyPure^®^RNA Kit (ER101-01, TransGen Biotech). RNA sequencing by an Illumina HiSeq instrument and data analysis were performed by Biomarker Technologies (Beijing, China). Illumina sequencing reads were mapped to reference genome *Glycine max* Wm82.a2.v1 (https://phytozome.jgi.doe.gov/pz/portal.htm).

### Statistical analysis

All experiments in this study were carried out at least three times, all of which showed similar results. The figures show only a representative result. Data in all bar graphs represent the mean ± SD. For digital statistical analysis, all statistical analyses were determined using SPSS software package. Asterisks indicate significant difference according to a Student’s *t*-test (\*\**P* < 0.01, * *P* < 0.05).

### Accession Numbers

Sequence data from this article can be found in Supplemental Information, and their corresponding identities are listed in Extended Data Table 1.

